# A Graph-Informed Modeling Framework Empowering Gene Pathway Discovery

**DOI:** 10.1101/2024.09.24.614661

**Authors:** Yihao Wang, Yue Wang, Jin Jin

**Affiliations:** Department of Biostatistics, Epidemiology and Informatics, University of Pennsylvania, Philadelphia, PA 19103, USA; Department of Biostatistics and Informatics, Colorado School of Public Health, Aurora, CO 80045, USA

**Keywords:** Block-coordinate descent, High-dimensional linear discriminant analysis, Gene regulatory networks, Gene expression, Mixed graphs, Gene pathway analysis

## Abstract

This study introduces a novel graph-informed modeling framework for improving the statistical analysis of gene expression data, particularly in the context of identifying differentially expressed gene pathways and gene expression-assisted disease classification in a high-dimensional data setting. By integrating gene regulatory network information into hypothesis testing for the difference between mean vectors and linear discriminant analysis, we aim to effectively capture and utilize previously validated external gene interaction information. Our method leverages a block-coordinate descent approach which enables us to incorporate mixed graph information into linear structural equation modeling, accommodating directed/undirected edges and potential cycles in gene regulatory networks. Extensive simulations under various data scenarios have demonstrated the effectiveness of our approach with improved power for gene pathway tests and disease classification over existing methods. An application to a lung cancer dataset from the Cancer Genome Atlas Program (TCGA) further exemplifies the potential of our graph-informed approach in empowering the detection of differentially expressed gene pathways and gene expression-based classification of different lung cancer stages. Our findings underscore the potential utility of incorporating gene regulatory network information in gene pathway analysis, setting the stage for future advancements in gene pathway discovery, disease diagnosis, and treatment strategies.

## Introduction

Gene discoveries, yielding numerous biologically significant insights on the genetic mechanism of human health conditions, have proven crucial for facilitating subsequent experiments and drug target identification^1–3^. The majority of statistical genetic discoveries have been relying on single gene analyses that reflect marginal associations rather than gene-gene interplay behind disease progression. On the other hand, gene pathways have proven to be successful drug targets for various diseases^4–9^, and pathway-level analysis aggregating signals from gene sets that function collectively could yield higher power and biologically more meaningful findings. Despite being a cornerstone for the evolution of disease etiology and targeted therapy, gene pathway prioritization is a methodologically under-investigated area. Furthermore, gene discoveries on both single gene or and pathway level face a critical issue of low power, partly due to the limited sample sizes. The expansive pathway databases have gathered rich gene regulatory network (GRN) information of genes and gene products across various biological processes^8,10–14^. Appropriately incorporating these priorly inferred GRN could greatly facilitate gene pathway analysis.

There has recently been a growing pursuit of statistical modeling approaches for effective gene pathway analysis. Several approaches have been proposed to incorporate gene interaction information within pathways, depicted through graphs as directed/undirected edges among genes (nodes), to compensate the typically inadequate sample size of gene expression datasets. For investigating differentially expressed gene pathways between populations or health conditions, a few high-dimensional two sample tests have been proposed utilizing gene interaction information. NetGSA test^15^ uses a linear mixed model to incorporate information on the magnitude of gene interactions, which, however, is often unknown or only partially known and thus hinders applicability of the approach. Another test named Graph T2 projects data onto a low-dimensional space spanned by top eigenvectors of the graph Laplacian of GRN and then performs dimension reduction^16^. While only requiring information on the presence/absence of gene interactions, it is likely to have compromised power when the signal is not coherent with the selected eigenvectors of the corresponding graph Laplacian. On the other hand, for broader applications beyond differential gene pathway analysis, a graph-guided Bayesian factor model was proposed to incorporate likely noisy GRN information and identify globally shared, partially shared and modality-specific latent factors in multi-modal genomic data^17^. It induces group structures in the effect of gene expression on health outcomes via gene interaction information, which proves effective for multi-omic data-based disease modeling.

The proposed graph-informed modeling framework improves inference in the high-dimensional setting through updating the precision matrix estimate by priorly inferred GRN information. The method only requires information on the presence/absence of interactions between genes. Built upon a previous framework named T2-DAG which can only handle directed acyclic graphs (DAG)^18^, our method could capture more complete graph information, allowing incorporation of mixed graphs without self-loop. The method utilizes structural equation modeling (SEM) to connect gene interaction with covariance matrix of gene expressions, which, while robust, typically can only handle DAG but not general mixed graphs featuring undirected edges and cycles. To address this challenge, we employ a block-coordinate descent (BCD) approach^19^ that enable SEMs to further handle mixed graph structures, capturing more comprehensive GRN information.

We implement the graph-informed framework for two purposes: hypothesis testing for differentially expressed gene pathways and classification of disease statuses based on gene expression levels. For hypothesis testing, we adopt the classical Hotelling’s T^2^ test framework but replace the sample precision matrix with an GRN-informed precision matrix estimate. For disease classification, we adapt the conventional Fisher’s Linear Discriminant Analysis (LDA) framework but replace the sample covariance matrix with the GRN-informed covariance estimate. Alternatively, besides the standard LDA framework, the idea can be applied to other classification algorithms, which we will show by simulations and real data examples. These two topics are of particular interest in cancer studies, where we aim to detect differentially expressed gene pathways and distinguish between cancers or different cancer stages based on gene expression levels.

We conduct extensive simulations under a variety of data settings, encompassing both low-dimensional and high-dimensional scenarios, where our graph-informed test shows good Type I error control and comparable or higher power compared to T2-DAG, which can only incorporate the largest sub-DAG within the graph. We further explored the classification capabilities of our graph-informed LDA framework. Simulation results suggest that integration of gene interaction information could improve classification accuracy substantially. We further demonstrate practicality of the method through a lung cancer application on data from the Cancer Genome Atlas Program (TCGA), where we detect differentially expressed gene pathways across different lung cancer stages as well as differentially expressed gene-based classification of lung cancer stages. Having demonstrated the promising performance of our GRN-informed approach over existing methods, our study sets the stage for future method development on unraveling the complex genetic mechanism of human health through integration of GRN information, which could assist development of advanced diagnostic tools and therapeutic strategies. Finally, the proposed GRN-informed modeling framework can be applied to a broader range of data types with available graph information, such as protein data analysis utilizing protein networks, brain functional magnetic resonance imaging (MRI) data analysis utilizing graphs describing structural connection in the brain, and modeling of microbiome data utilizing phylogenetic tree information.

## Method

Our method comprises a graph-informed modeling framework followed by two distinct applications: hypothesis testing for differentially expressed gene pathways and gene expression-assisted disease classification. The graph-informed approach builds upon its predecessor given in T2-DAG^18^, which was limited to incorporating only the directed acyclic components of a graph. This novel method extends to also including undirected edges and cycles, thus incorporating nearly the entire graph structure except for self-loops. Suppose we have the adjacency matrix, *A* = (*a*_*jk*_) ∈ ℝ^*p*×*p*^ available from some pathway databases, where *a*_*jk*_ = 1 indicates the presence of the edge *k* → *j*, i.e., node *k* affecting node *j*, and *a*_*jk*_ = 0 otherwise, in a possibly mixed graph *G*. Here, the nodes represent individual genes while the edges represent the interactions between them, encompassing both activation and inhibition dynamics. If *a*_*jk*_ = 1 and *a*_*kj*_ = 1, then we say the edge between nodes *j* and *k* is undirected or bidirectional. To proceed with structural equation modeling utilizing this adjacency matrix, we need to further assume sparsity of *A* and that there is no self-loop in *G*, i.e. *a*_*ii*_ = 0, *i* ∈ [*p*]. Detailed assumptions are described in Drton, Fox and Wang (2019)^19^.

While SEM offers a framework for integrating network information in the estimation of covariance matrix^18^, it faces limitations when encountering graphs that extend beyond DAGs. For example, when interactions include undirected edges or cycles among nodes, the computation of maximum likelihood estimates (MLEs) in linear SEMs with Gaussian errors in the structural equations become challenging, where solutions to standard regressions are unidentifiable and the general quasi-Newton methods for nonlinear optimization face convergence issues and require careful choice of initial values^20^. To overcome these limitations, innovative improvements on SEM, such as the BCD method proposed by Drton, Fox, and Wang (2019)^19^, have been introduced. BCD is particularly appealing for its ability to avoid the need for selecting an ideal initial value and the subsequent convergence issues. Its closed-form solutions for each block-update problem in linear

SEM align well with the challenges presented by our consideration of incorporating mixed graph information into a linear SEM framework. BCD operates by iteratively updating parameters related to each variable (block) while maintaining all other parameters constant until the log-likelihood function’s partial maximization reaches at a pre-specified tolerance, thereby ensuring convergence to a solution that accounts for the complexity of interaction structures in mixed graphs.

Given a set of centered, unscaled, and normally distributed random variables *Z* = (*Z*_1_, *Z*_2_, … , *Z*_*p*_) , we consider a set of SEMs, *Z* = *ZB* + *ε* , where *ε* ∼ *N*(0, Ω) is a set of independent (i.e., Ω = diag(*ω*_1_, *ω*_2_, … , *ω*_*p*_)), normally distributed error terms. We can then derive the covariance matrix of *Z* in the form of *S* = (*I* − *B*^*T*^)^−1^Ω(*I* − *B*)^−1^ and the precision matrix in the form of *S*^−1^ = (*I* − *B*)Ω^−1^(*I* − *B*)^*T*^ . By treating directed edges and undirected edges separately but simultaneously in each block update, the BCD algorithm dismantles the sparse adjacency matrix *A* into two matrices: *B* = (*β*_*ij*_) for directed edges, sometimes referred to as structural parameters, and Ω = (*ω*_*ij*_) for undirected edges, often used to represent the latent variables that are potentially the common causes of both directed and undirected effects. Thus, the notations *B* and Ω here also conform with those in the SEM framework after BCD updates. For computational convenience, we also manually set the diagonal entries of Ω to 1, i.e. *ω*_*i*_ = 1, *i* = 1, 2, … , *p*. Then, at convergence, the algorithm returns the updated matrices, *B̂*_*BCD*_ and Ω̂_*BCD*_, and with these BCD updated estimates, the covariance matrix of gene expressions, which is consistently nonsingular, and the precision matrix can be reconstructed, which we refer to as the GRN-informed covariance matrix estimate *Ŝ* and precision matrix estimate *Ŝ*^−1^.

This GRN-informed modeling framework is particularly useful in the high-dimensional setting with *p* > *n*_1_ + *n*_2_, where *n*_1_ and *n*_2_ are the sample size for two different conditions, in which case the classical Hotelling’s T^2^ test and LDA algorithm become inapplicable due to the ill-defined sample covariance matrix. We first look at the first application of our GRN-informed approach, which is to test for the difference in the mean values of gene expression levels between two conditions (denoted by *X*^(1)^ and *X*^(2)^), formally, to test the hypothesis *H*_0_: ***μ*_1_** = ***μ*_2_** against *H*_*a*_: ***μ*_1_** ≠ ***μ*_2_**. Although proven to be the uniformly most powerful invariant (UMPI) test^21^, Hotelling’s *T*^2^ test could be mistaken if the precision matrix is falsely imposed, especially when the covariance matrix is singular under high-dimensional scenarios. Thus, by integrating the edge information in GRN into the analysis and obtaining the precision matrix estimate with BCD- estimated SEM coefficients, we could replace the ill-proposed sample precision matrix in the

Hotelling’s *T*^2^ test statistic by the GRN-informed precision matrix *Ŝ*^−1^ and obtain a test statistic 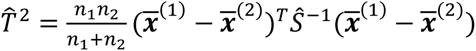. When a comprehensive set of pathways are being analyzed, the test will assess the significance of each individual pathway in terms of differential expression between two health conditions. By pinpointing the significant expressed pathways and their gene interaction features, this hypothesis testing procedure effectively serves as an initial feature engineering step to identify the differentially expressed gene pathways, laying the groundwork for various downstream analyses.

The GRN-informed approach also leads to a second application of distinguishing between different disease statuses based on differential gene expression patterns across pathways. LDA is a commonly implemented framework for such classification tasks, with a primary goal of maximizing separability between predefined clusters. In the context of this study, LDA classifies observations into distinct lung cancer stages based on gene expression patterns. Fisher’s Linear Discriminant Analysis, which is the commonly used LDA framework, is recognized for its asymptotic optimality in scenarios characterized by a fixed and low dimension of features (*p*), as illustrated in numerous studies^22,23^. LDA distinguishes between different clusters by identifying a linear axis (or multiple axes for multi-class problems) that most effectively separates between classes. Through data projection onto these axes, LDA reduces the dimension of the data while preserving key class-discriminatory information. The essence of the approach is the derivation of an optimal projection vector *v* from the equation *y* = *v*^*T*^*x*, aiming to maximize between-class variability while minimizing within-class variability.

LDA assumes a normal data distribution and homogeneous covariance matrices across different classes. In the binary classification scenario, it computes a total scatter matrix *S*_*ω*_ = *n*_1_Σ̂_1_ + *n*_2_Σ̂_2_, which is a weighted sum of the sample covariance matrices for the two groups. It then seeks solution *ω*^∗^, the optimal projection vector, for the equation *S*_*ω*_*ω*^∗^ = ***μ***_2_ − ***μ***_1_. Classification is then performed by comparing the projection of observations on *ω*^∗^ against the project of group means, then classifying observations to the nearest group. In a more conventional form of Fisher’s linear discriminant rule, suppose a new observation is being classified based on the observed data feature ***Z*** within the parameter space *θ* = (***μ***_1_, ***μ***_2_, Σ), the classifier is given by

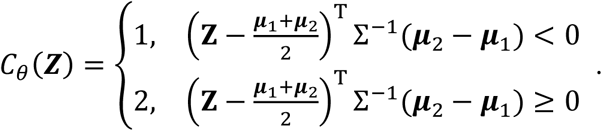

In the high-dimensional setting where the number of features (*p*) is larger than the number of observations (*n*), Fisher’s discriminant rule fails due to singularity of the covariance matrix estimate (Σ̂). By replacing the ill-posed precision matrix (Σ̂^−1^) by a graph-informed estimate, our proposed approach offers an effective solution to this inherent singularity problem in the general LDA framework. It differs from the commonly implemented remedial approach of adding a small value to the diagonal entries of the singular covariance matrix, an ad hoc approach to forcing the covariance matrix to be invertible. Following the implementation of this graph-informed precision matrix estimate, our approach then calculates the posterior probability of the observation belonging to each class using the Bayes’ theorem as in the conventional LDA framework.

### Simulation Studies

#### Simulation Settings

We conduct extensive simulations to assess the performance of our GRN-informed approach in terms of testing and classification performance. For hypothesis testing, we refer to our method as “T2-MG”, which employs a mixed graph (MG)-informed *χ*^2^ or *Z* test statistic with asymptotic validity under certain conditions^18^. We considered various settings of sample sizes (*n*_1_ = *n*_2_ ), number of genes in a pathway (*p*), descendant/children nodes in the graph (genes that are affected by at least one other gene in a pathway, *p*_0_), number of differentially expressed genes (*q* = 0.5*p*), and a maximum number of directed edges toward one gene ( *d* ). We investigated method performance in 5 different settings of *n*_1_ , *n*_2_ , and *p* under both low-dimensional and high-dimensional scenarios, and in each setting, we set the number of parent genes to either *p*_0_ = 0.4*p* or *p*_0_ = 0.8*p*.

We simulate data by first generating the adjacency matrix for a sparse DAG given *p*_0_ and *d*, where *d* follows a multivariate negative binomial distribution with *p*_0_ observations and a success probability of 0.6 for *p*_0_ = 0.4*p* or 0.8 for *p*_0_ = 0.8*p*. Then, without loss of generality, we introduce a loop between the first three nodes (C to B, B to A, and A back to C) as a simple representation of a mixed graph characteristic. We then converted a random half of the remaining directed edges into undirected (bidirectional) edges. This process may introduce additional loops beyond the one that is initially added, which involves nodes A, B, and C. The resulting adjacency matrix, *A* = (*a*_*jk*_) ∈ ℝ^*p*×*p*^, now reflects the underlying gene-gene interaction network and is used for further data generation. As previously mentioned, when implementing the BCD algorithm, it is essential to separate directed and undirected edges from the adjacency matrix *A*. While this separation might seem redundant to implement here as we already start with generating a DAG, it is yet crucial due to the randomness introduced by converting some directed edges into undirected ones, which can formulate unknown loops. Therefore, we conduct the separation after the adjacency matrix *A* is generated. This step divides adjacency matrix *A* into *B*, the matrix for directed edges, and Ω, the matrix for undirected edges with all diagonal elements set to 1. Note that the BCD algorithm cannot process self-loops, hence all diagonal entries in *A* are 0. After generating the adjacency matrix, we proceed to simulate observed data for the two groups, *X*^(1)^and *X*^(2)^ . We adopt the data generation protocol in T2-DAG^18^, initiating with the true data coefficient matrix *Q* and the true diagonal error variance matrix *R* = diag(0.2) in the SEM ***X*** = ***X****Q* + ***ε***, where ***ε*** ∼ *N*(**0**, *R*). Starting with the raw *X*^(1)^ and *X*^(2)^, both having zero mean, we maintain the mean of *X*^(1)^ at 0 while assigning a distinct value, denoted by the signal strength *δ*, to the mean of *X*^(2)^, thereby establishing the population means ***μ*_1_** = 0 and ***μ*_2_** = *δ*. We then merge the two sets of data *X*^(1)^ and *X*^(2)^ to obtain the full data, which we denote by *Y*.

We conducted hypothesis tests based on graph-informed Z statistic (*T*_*Z*_) or Chi-squared test statistic 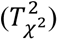 using either the largest sub-DAG (T2-DAG test^18^) or the complete mixed graph information excluding possible self-loops (the proposed test). The structure of the adjacency matrices is distinct between the DAG and MG setups when fitting the linear SEM. In the MG case, we employ the full adjacency matrix with the BCD algorithm to estimate graph-informed covariance and precision matrices. In contrast, in the DAG case, the largest subset of *A* that captures only the directed and acyclic edges is used, and the classical SEM fitting with linear regressions is used.

Based on the graph-informed covariance matrix estimates *Ŝ*_*DAG*_ (T2-DAG^18^) or *Ŝ*_*MG*_ (the proposed method), we then compute the corresponding test statistics *T*_*DAG*−*Z*_, 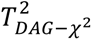 (T2-DAG tests^18^), *T*_*MG*−*Z*_, and 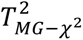 (the proposed tests). Our objective is to assess the Type I error rates (when *δ* = 0 , indicating no true mean differences between *X*^(1)^ and *X*^(2)^ groups) at a significance level of *α* = 0.05, as well as the statistical power (when |*δ*| > 0). Considering that the T2-DAG test (using either *T*_*DAG*−*Z*_ or 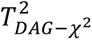) has already been shown to outperform other existing methods with a higher power and a well-controlled Type I error rate^18^, our study simplifies the comparison by focusing solely on the proposed MG-informed tests against T2-DAG tests.

The simulation settings for LDA-based classification closely mirror those for hypothesis testing, but with unequal sample sizes *n*_1_ and *n*_2_ to reflect the unbalanced sample scenario. To better illustrate the AUC differences between the two methods, we have shifted from using customized signal strengths for different settings to a unified sequence of signal strengths starting from 0 and increasing by 0.05 up to 0.25, at which point we anticipate that the LDA classifications reach an AUC value close to 1, assuming *X*^(2)^ as the case group and *X*^(1)^ as the control. In addition to DAG-LDA and MG-LDA, we also implement the classical LDA^24,25^ as a benchmark for comparison. We purposely do not address the issue of singularity in high-dimensional settings for the Benchmark LDA, provided that it does not result in computational errors. This approach offers a direct comparison of the Benchmark LDA with MG-LDA and DAG-LDA when confronted with the challenges of high-dimensionality. Classification performance is evaluated based on AUC estimated from Leave-One-Out Cross Validation (LOOCV).

### Simulation Results

The hypothesis testing results are summarized across 500 replicates per simulation setting (Figure 1, Figures S1 – S3). Given the inadequacies in asymptotic properties of the BCD inference, we considered implementing *T*_*MG*−*Z*_ in a permutation test as an alternative. We observe a consistent pattern across different simulation settings that *T*_*MG*−*Z*_ outperforms *T*_*DAG*−*Z*_ with a higher power and a slightly increased but still relatively well controlled type I error rate. while 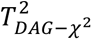 and 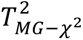 have similar performance. Meanwhile, 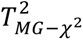 exhibits marginal improvements in power and a similar type I error control compared to 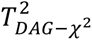. Furthermore, as *p*_0_, the number of descendant genes, increases, T2-MG exhibits more noticeable advantage over T2-DAG in terms of testing power for both *Z* and Chi-squared tests. These results suggest that MG-informed tests, especially the Z tests, can better uncover differential gene expression patterns by introducing information on gene interactions from external pathway databases. In the context of disease diagnostics, the higher power of the *T*_*MG*−*Z*_ test is often deemed more valuable. This approach favors a more aggressive testing strategy, accepting a higher risk of false positives for the diseased group over the control group, while maintaining a higher sensitivity at detecting the true diseased group. Such a strategy is preferable in scenarios where missing a true positive could have more severe consequences than falsely identifying a negative case as positive. Finally, we have previously shown that T2-DAG tests are highly robust to GRN misspecification (40% missing or redundant edges)^18^, and thus we anticipate a similar robustness of T2-MG tests.

**Figure 1:**
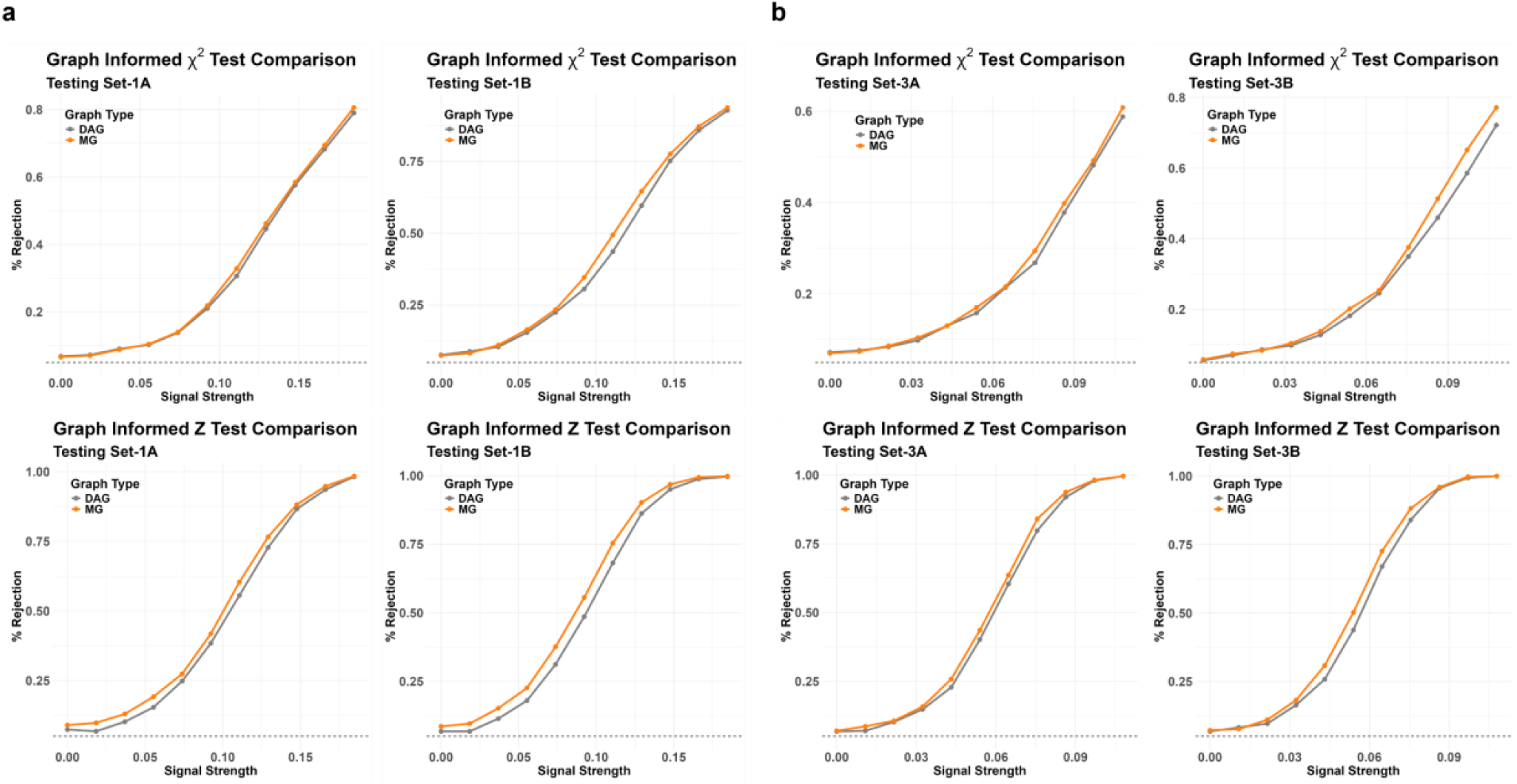
Simulation results for hypothesis testing of differentially expressed gene pathways. Type I error rates (signal strength *δ* = 0) and statistical powers (|*δ*| > 0) are compared based on graph-informed *Z* (row 1) and *χ*^2^ (row 2) type tests between T2-DAG tests utilizing the largest sub-DAGs and the proposed tests utilizing complete MG information. Simulations were conducted with either setting A: *p*_0_ = 0.4*p* (columns 1 and 3) or setting B: *p*_0_ = 0.8*p* (columns 2 and 4). Results under other simulation settings are summarized in Figures S1 – S3. **a.** Simulation setting 1 (1A and 1B) highlights low-dimensional scenarios with equal sample sizes (*n*_1_ = *n*_2_ = 50) and a modest number of genes within a pathway (*p* = 20). **b.** Setting 3 (3A and 3B) highlights the high-dimensional scenarios, with sample sizes *n*_1_ = *n*_2_ = 50 and a larger number of genes within each pathway (*p* = 100). Comparison of DAG-based tests with other existing tests is summarized in Jin and Wang (2022)^18^.

For classification, we implement and compare a series of different classification approaches, including MG-LDA and DAG-LDA, our proposed graph-informed LDA algorithm utilizing complete graph information (excluding self-loops) and the largest sub-DAG, respectively, Benchmark LDA: the standard Fisher’s LDA, Penalized LDA^26^, and Penalized logistic regression (i.e., Penalized GLM)^27^. To illustrate the broad utility of our graph-informed approach, we additionally consider AdaLDA^22^, a recent LDA algorithm specifically designed for high- dimensional settings with a data-driven and tuning-free classification rule based on an adaptive constrained *L*_1_-minimization approach, and implement AdaLDA-GRN, which incorporates the GRN-informed covariance matrix generated by our GRN-informed method into the AdaLDA framework.

Generally, in the relatively low dimensional settings (Figure 2, Figures S4 – S6, where *n*_1_ = *n*_2_ =50 and *p* = 20 or 40), the classification accuracy does not have a big difference between different approaches. But in the relatively high dimensional settings (Figures S7 – S10, where *n*_1_ = *n*_2_ = 100 and *p* = 100 or 300), we observe a much bigger difference in AUC across different dimension reduction approaches. Overall, both MG-LDA and DAG-LDA demonstrate superior performances in classifying binary outcomes when compared to Benchmark LDA, especially when the differences between the two classes are large (Figure 2, Figures S4 – S10). The distinction in performance is less obvious between MG-LDA and DAG-LDA. Compared to DAG-LDA, MG-LDA tends to exhibit slightly higher AUC in all scenarios, especially in the high-dimensional scenarios with sample sizes. On the other hand, DAG-LDA exhibits stable performance, showing no decreases and, in rare cases, marginal increases in AUC in the low-dimensional settings or when the between-class difference is very large, in which case the information on the presence/absence of gene interactions becomes less informative for classification. In all other cases, the performance between MG-LDA and DAG-LDA does not show noticeable differences. Compared to the classical dimension reduction technique (penalized LDA and penalized GLM), further utilizing GRN information by our proposed graph-informed classifiers can lead to additional improvement. Our graph-informed modeling framework also show its effectiveness when applying to other LDA framework, such as AdaLDA, where it can substantially improve the AUC especially when there is a relatively large heterogeneity between the two populations/conditions (Figure 2, Figures S4 – S10, when *δ* ≥ 0.10).

**Figure 2:**
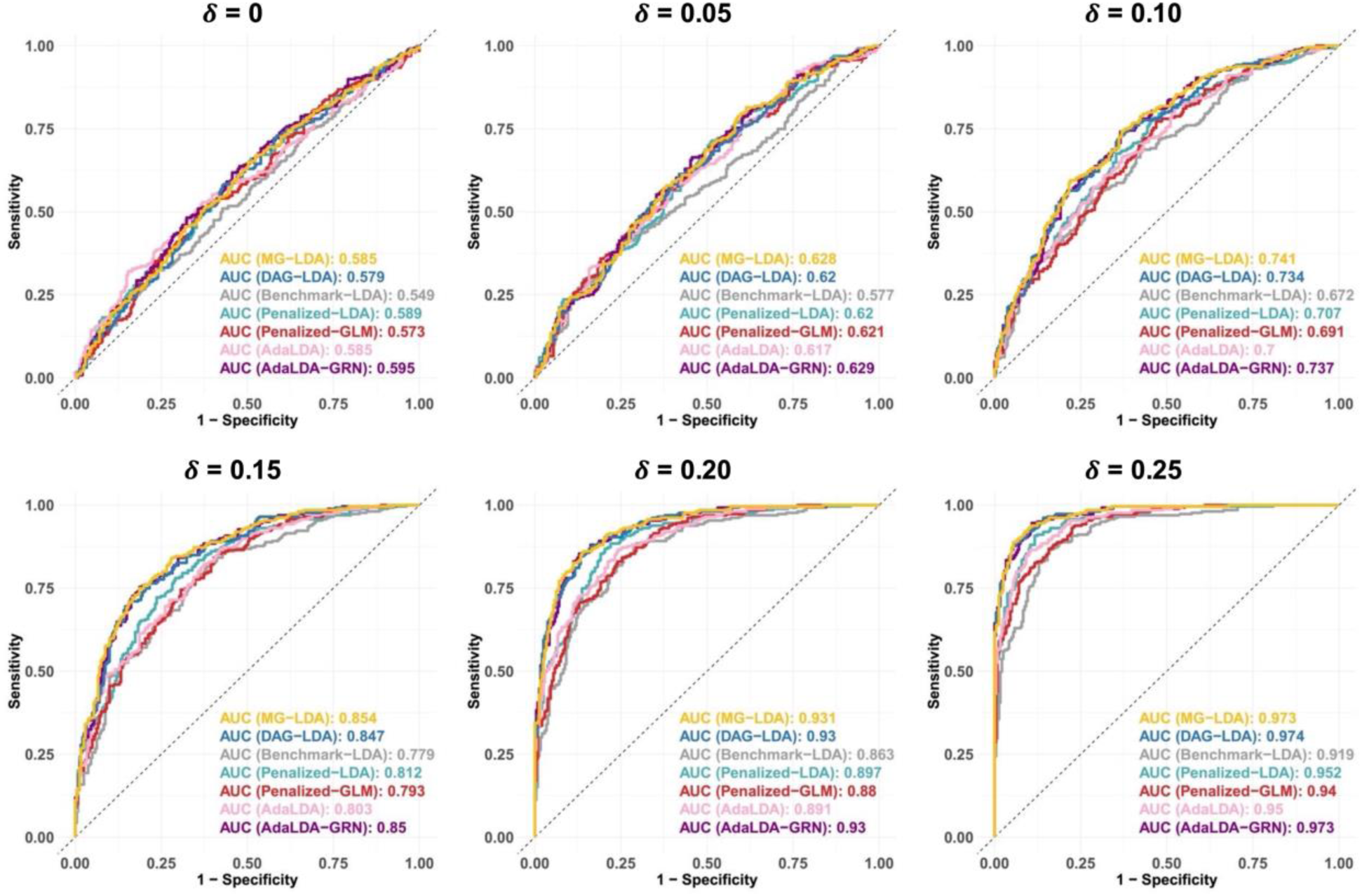
Simulation results showing the classification performance of various LDA methodologies. We consider MG-LDA and DAG-LDA: the proposed graph-informed LDA algorithm utilizing complete graph information (excluding self-loops) and the largest sub-DAG, respectively; Benchmark LDA: the standard Fisher’s LDA; Penalized LDA, Penalized GLM, AdaLDA, and AdaLDA-GRN, which incorporates the GRN-informed covariance matrix generated by our GRN-informed method into the AdaLDA framework. We showcase simulations results given *n*_1_ = *n*_2_ = 50, *p* = 40, and *p*_0_ = 0.4*p* under various settings of the mean difference between two groups *δ* = 0, 0.05, … , 0.25 . Results under other simulation settings are summarized in Figures S4 – S10.

**Figure 3:**
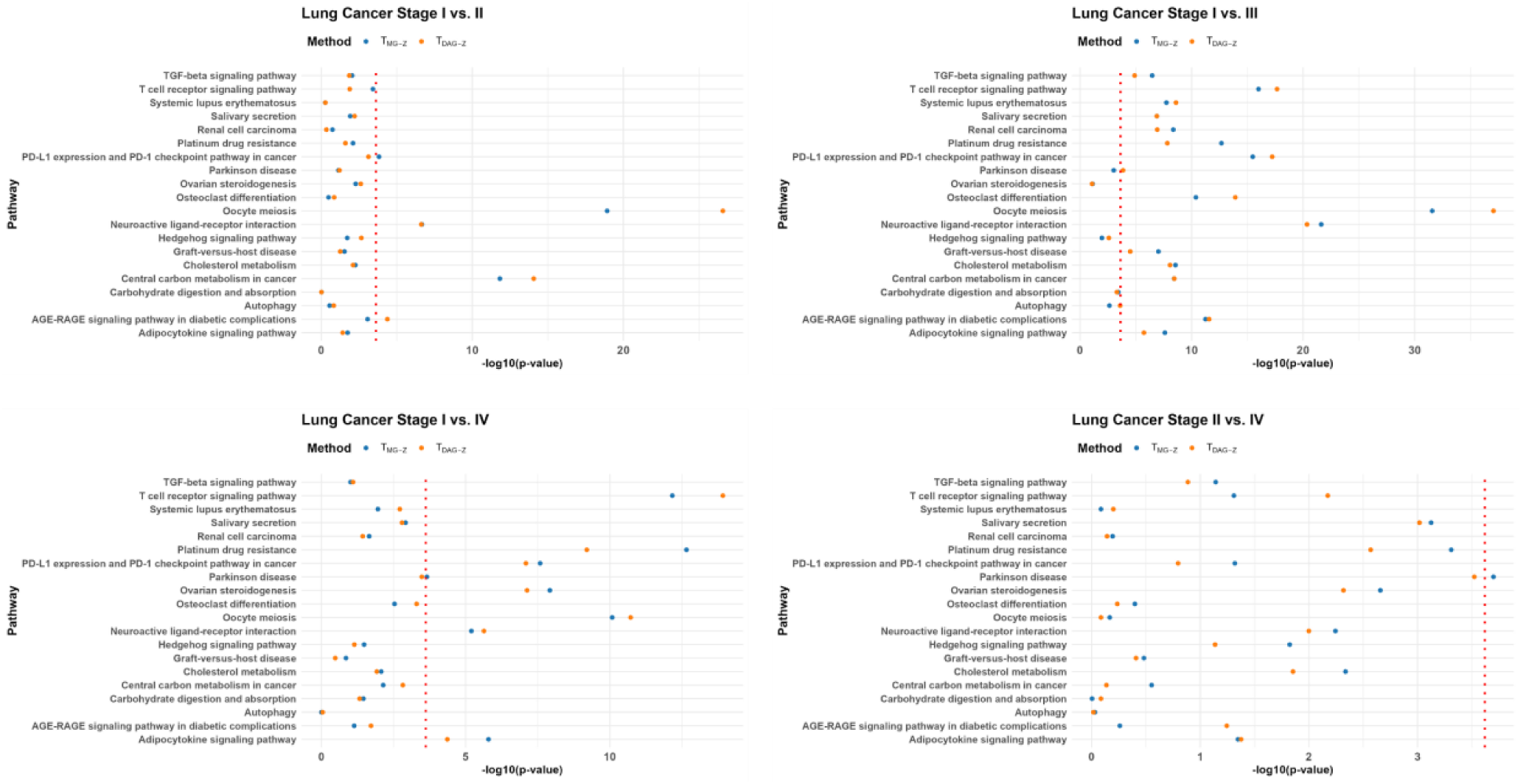
Results of the hypothesis tests based on *T_MG−Z_* and *T_DAG−Z_* for 20 randomly selected KEGG pathways across different lung cancer stages. The red dashed line within each graph marks the Bonferroni adjusted significance threshold, *α* = 0.05/208. Pathways whose transformed p-values are on the right-hand side of the line are identified to be significantly differentially expressed between the corresponding pair of lung cancer stages.

### Gene expression analysis across lung cancer stages with TCGA data

We apply the tests and classifiers to a dataset comprising gene expression profiles of lung cancer patients from TCGA^28,29^. The data used in our study was preprocessed so that non-expressive values were normalized to the minimum observed expression value across the dataset, followed by a base-2 logarithmic transformation to approximate normal distribution for all lung cancer stages, a methodology consistent with the approach outlined by Cai and Jiang^30^. The dataset featured gene expression levels for 20,429 genes within lung tissue samples from 513 patients, stratified by lung cancer stages I through IV (*N*_1_ = 278, *N*_2_ = 124, *N*_3_ = 84, and *N*_4_ = 27).

Additionally, 59 normal tissue samples were included from the 59 of the lung cancer patients, which are excluded from the MG-informed tests and classifications. *H* = 208 human pathways were considered, which reflect biological functions related to signaling, metabolic processes, and human diseases, as cataloged in the KEGG database^31,32^. As the KEGG and other pathway databases undergo continuous expansion, the breadth of data they collects has notably increased^8,10–14^, reflecting the ongoing development of the pathway databases as comprehensive resources for GRN informed pathway analysis.

The first part of our graph-informed analysis involves hypothesis testing for identifying pathways that have differentially expressed genes between lung cancer stages. We use the same hypothesis testing procedures as the one described in the simulation studies. For the comparison between each pair of lung cancer stages, we apply *T*_*MG*−*Z*_ and *T*_*DAG*−*Z*_ to each pathway and apply a Bonferroni correction to the significance threshold, setting *α* = 0.05/*H* with *H*=208. The Chi-squared tests had similar performance and are not summarized here. Besides identifying differentially expressed pathways across lung cancer stages, this hypothesis testing step effectively serves as a primary feature engineering step that could drastically reduce the number of genes from 20,429 to a more manageable number for downstream analyses.

Among the six different pairs of lung cancer stages, four have significant pathways identified (Stage I vs. Stage II: 54 pathways; Stage I vs. Stage III: 149 pathways; Stage I vs. Stage IV: 81 pathways; Stage II vs. Stage IV: 14 pathways; based on *T*_*MG*−*Z*_ ). Our analyses thus only focus on these four pairs of lung cancer stages. Given the high consistency in classification performance of the DAG-informed and MG-informed classifiers in simulation studies, we consider choosing only one of the two graph-informed classifiers. We observe that the process of extracting the largest sub-DAG from the full graph can be extremely time consuming when *p* > 150 and often encounter failures, and thus we propose to only use the MG-informed LDA approach for our analysis that utilize complete graph information.

Upon identifying significant pathways between two lung cancer stages, a subset of differentially expressed genes is obtained, which will serve as the input for gene expression-based LDA classification. One thing to note is that, while gene expression data alone may not have sufficient power for classifying disease statuses, it can be combined with other data modalities to develop classifiers that can reach a high enough classification accuracy to be implemented in real applications. The graph-informed LDA framework we present here is used to demonstrate the effectiveness of incorporating GRN information in facilitating the investigation of disease-related differential gene expression patterns, but not necessarily to generate classifiers for clinical use. When performing classification, we aggregate signals of differential expression across all significant pathways identified in the hypothesis testing step, with a large 0-1 adjacency matrix *A* summarizing presence/absence of gene interactions across all pathways. Consequently, the graph-informed covariance matrix is estimated by a single BCD process.

Figure 4 shows the ROC curves of classification of lung cancer stages using MG-LDA, Benchmark LDA, Penalized LDA, Penalized GLM, AdaLDA, and AdaLDA-GRN. MG-LDA has a higher AUC than Benchmark LDA in all four comparisons, suggesting that our proposed MG-LDA approach can improve the discriminatory ability of gene expression-based classifiers. Specifically, the Stage I vs. Stage IV comparison shows a more notable difference in AUC between classifiers, with the MG-LDA achieving AUC=0.658, which is much higher than that of Benchmark LDA (AUC=0.537) and penalized GLM (AUC=0.601). AdaLDA-GRN, which accommodates our proposed graph-informed covariance matrix estimate, also show promising performance with a similar or higher AUC than the original AdaLDA algorithm.

**Figure 4:**
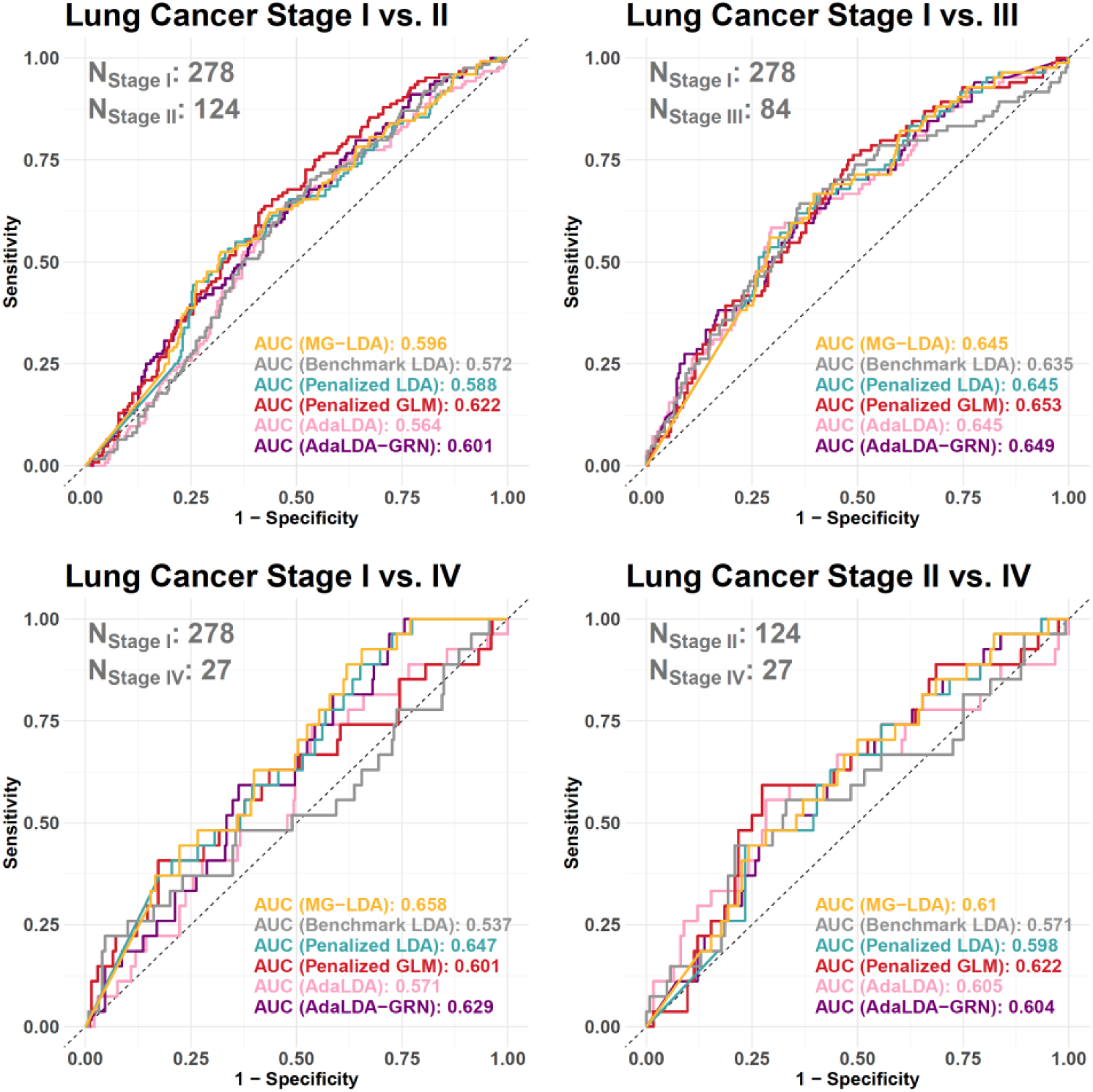
Classification results based on various approaches across four pairs of lung cancer stages. The numbers of genes utilized, i.e., the total number of genes in the differentially expressed pathways identified by our hypothesis tests for the corresponding lung cancer stages, are: 1862 (I vs. II), 2857 (I vs. III), 2036 (I vs. IV), and 749 (II vs. IV), respectively. No differentially expressed pathway is identified between lung cancer stages II vs. III and III vs. IV, and thus classification was not conducted for these two pairs.

Notably, none of the AUC values reach or exceed the threshold of 0.7. Therefore, while MG-LDA shows much promise in improving the gene expression-based classification of lung cancer stages by incorporating GRN information, classifiers for lung cancer stages using only gene expression information are generally not powerful enough to be implemented in real practice, and additional data modalities are required to train more powerful classifiers. Even so, the results here highlight the potential of the proposed graph-informed modeling approach in improving the power of gene pathway analysis.

## Discussion

We propose a novel approach to integrating external pathway information on gene interactions into pathway analysis, specifically, the detection of differentially expressed gene pathways and gene expression-based classification across health conditions. We utilize a BCD algorithm that allows our method to incorporate edge information from mixed graphs, i.e., absence/presence of directed or undirected gene interactions, under a linear SEM framework. The method is particularly appealing in the high dimensional settings, where inference can be effectively improved by replacing the singular sample covariance matrix by a GRN-informed covariance estimate. The effectiveness and robustness of our method has been rigorously validated through extensive simulations in both low-dimensional and high-dimensional settings, as well as an application to analyze differences between different stages of lung cancer based on a dataset from TCGA. While gene expression data may not have sufficient power for classifying between different lung cancer stages in the TCGA data example, incorporating GRN information can still lead to substantial improvement in classification accuracy, which shows the great potential of our graph-informed approach in improving the power of gene pathway analysis.

The proposed graph-informed modeling approach also has limitations. A key challenge lies in the computational demand, particularly for implementing the BCD algorithm to solve linear SEM with mixed graphs. Although incorporating mixed graphs makes our algorithm slower, its computational speed is still acceptable for the typically gene pathway analysis. It also tends to show slightly better performance than incorporating only the directed acyclic components of the graphs. Furthermore, incorporating only the directed acyclic components of a graph require us to extract the largest sub-DAG from the graph, which can become increasingly time-intensive when *p* > 3000. We thus recommend the MG-informed approach over DAG-informed approach in the general applications of our method.

In the data example where we classify between different lung cancer stages based solely on individuals’ gene expression profile, while MG-LDA shows the potential to fundamentally improve the classification power of Benchmark LDA, the AUC remains limited and does not exceed 0.7. This suggests that gene expression data alone may not be sufficient for classification of many health conditions, including lung cancer stages. For these specific diseases and conditions, we may need to further collect data on other risk factors, such as lifestyle factors and environmental factors, to achieve a desirable classification accuracy. Still, our results justify the usefulness of incorporating previously validated external GRN information in improving the power of gene expression analysis, providing a novel modeling framework to the literature of statistical genomic research.

Looking ahead, there are also extensive opportunities to extend our graph-informed modeling framework to more general data applications. For example, the current LDA framework can be extended to the Quadratic Discriminant Analysis (QDA) framework with varying covariance matrices between different groups, given that the assumption of equal covariance in LDA might not always be met. The showcased binary classification framework can also be naturally extended to classify more than two groups^33,34^. Furthermore, we anticipate the possibility that future studies utilize advanced machine learning and AI technologies, such as Graph Neural Networks (GNN)^35^ and Transformer Neural Networks (TNN)^36^, to further advance gene pathway discovery.

While we initially developed our method based on the gene expression data-based applications, the general graph-informed modeling framework we propose, including the graph-informed hypothesis test and classification algorithm, can be applied to a wider range of applications with a variety of data modalities. Examples include protein data analysis utilizing protein networks^37–39^, where protein data has proven powerful for analyzing genetic mechanisms of various traits and disease outcomes^40–42^; and multi-omic data integration for gene pathway discovery and disease risk prediction^43^ utilizing available multi-omic GRN information on the bulk tissue level^44–46^ or single cell level^47–54^. Other examples include the analysis of functional magnetic resonance imaging (MRI) data combined with external structural connectivity information^55,56^ and microbiome data analysis incorporating phylogenetic tree information^57,58^. Essentially, any data application that can be represented as analyzing observed data of a group of nodes with information available on the absence/presence of edges can potentially benefit from our proposed graph-informed approach.

In summary, the proposed graph-informed method presents a significant step in facilitating statistical analysis of gene pathways. While the method has promising performance in hypothesis testing and classification through simulation studies and an application to gene expression-based lung cancer study, further research is demanded to broaden its utility in applications to other human conditions, other analytical tasks (e.g., multi-class disease classification, analysis of continuous health conditions under a regression framework), and other data types (e.g., protein data, brain MRI data, microbiome data). As the graph-informed method is being continuingly refined and extended, it could hold the potential to become a widely implemented tool in human health research and beyond.

## Code Availability

The code for conducting GRN-informed gene pathway test and disease classification can be accessed at https://github.com/Jin93/Graph-Informed-Model.

## Acknowledgements

This work was supported by the National Institutes of Health (NIH) grants R00 HG012223 (J.J.).

