## Supplementary Figures for "A Graph-Informed Modeling Framework Empowering Gene Pathway Discovery"

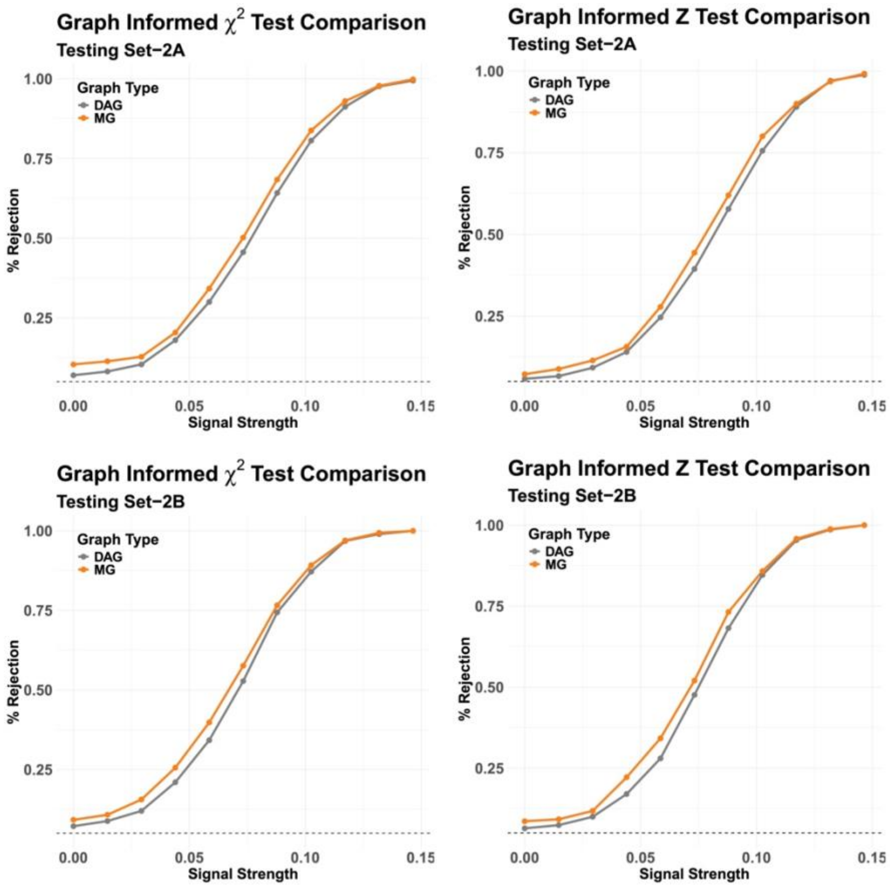


**Figure S1: Simulation results for hypothesis testing of differentially expressed gene pathways given $n_1 = n_2 = 50$ and $p = 40$.** Type I error rates (signal strength $\delta = 0$) and statistical powers ($|\delta| > 0$) are compared between either the graph-informed $\chi^2$ test (column 1) or the graph-informed $Z$ test (column 2) and T2-DAG tests utilizing the largest sub-DAGs and the proposed tests utilizing complete MG information. Simulations were conducted with either setting A: $p_0 = 0.4p$ (top panel) or setting B: $p_0 = 0.8p$ (bottom panel).

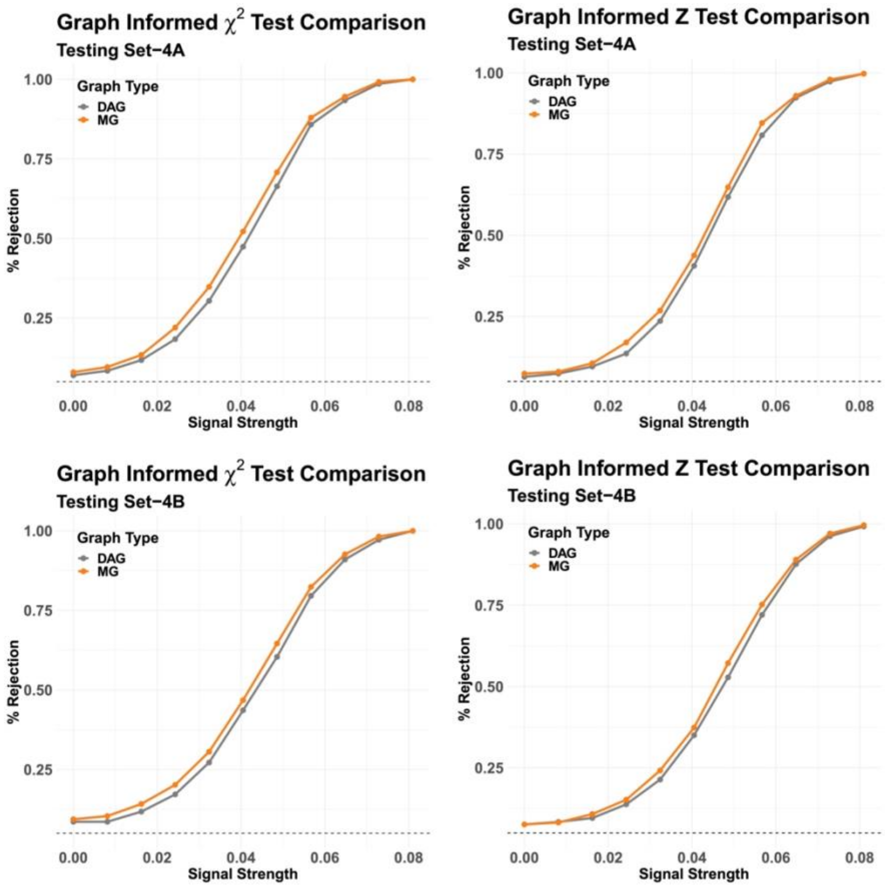


**Figure S2: Simulation results for hypothesis testing of differentially expressed gene pathways given $\boldsymbol{n_1 = n_2 = 100}$ and $\boldsymbol{p = 100}$.** Type I error rates (signal strength $\delta = 0$) and statistical powers ($|\delta| > 0$) are compared between either the graph-informed $\chi^2$ test (column 1) or the graph-informed $Z$ test (column 2) and T2-DAG tests utilizing the largest sub-DAGs and the proposed tests utilizing complete MG information. Simulations were conducted with either setting A: $p_0 = 0.4p$ (top panel) or setting B: $p_0 = 0.8p$ (bottom panel).

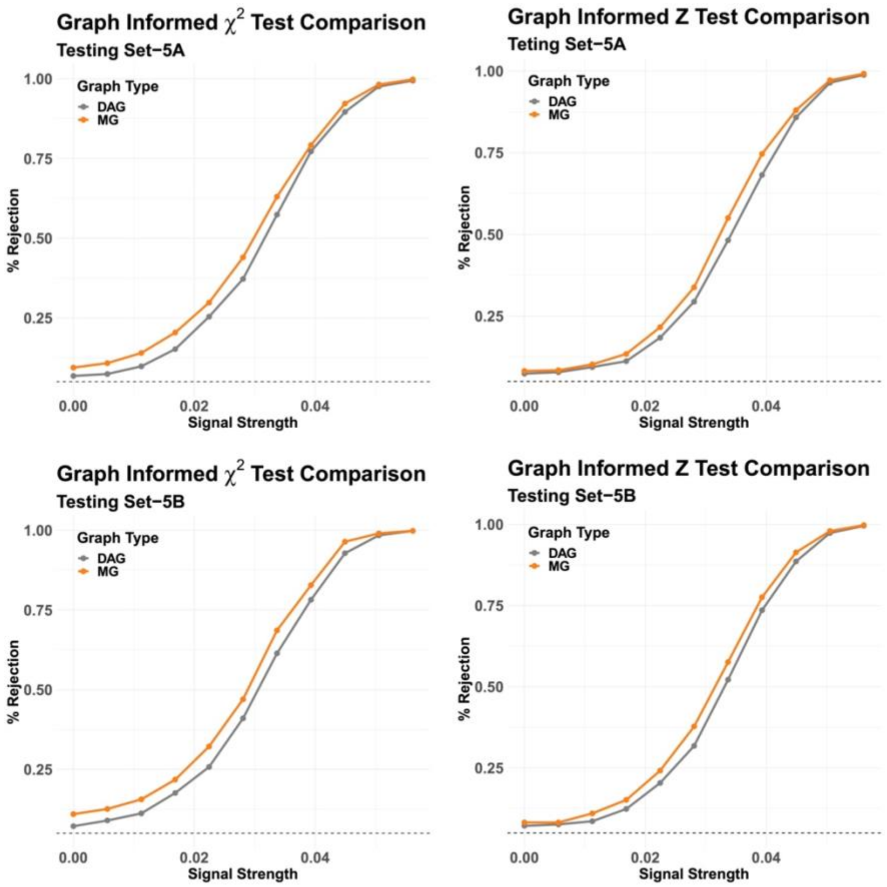


**Figure S3: Simulation results for hypothesis testing of differentially expressed gene pathways given $\boldsymbol{n_1 = n_2 = 100}$ and $\boldsymbol{p = 300}$.** Type I error rates (signal strength $\delta = 0$) and statistical powers ($|\delta| > 0$) are compared between either the graph-informed $\chi^2$ test (column 1) or the graph-informed $Z$ test (column 2) and T2-DAG tests utilizing the largest sub-DAGs and the proposed tests utilizing complete MG information. Simulations were conducted with either setting A: $p_0 = 0.4p$ (top panel) or setting B: $p_0 = 0.8p$ (bottom panel).

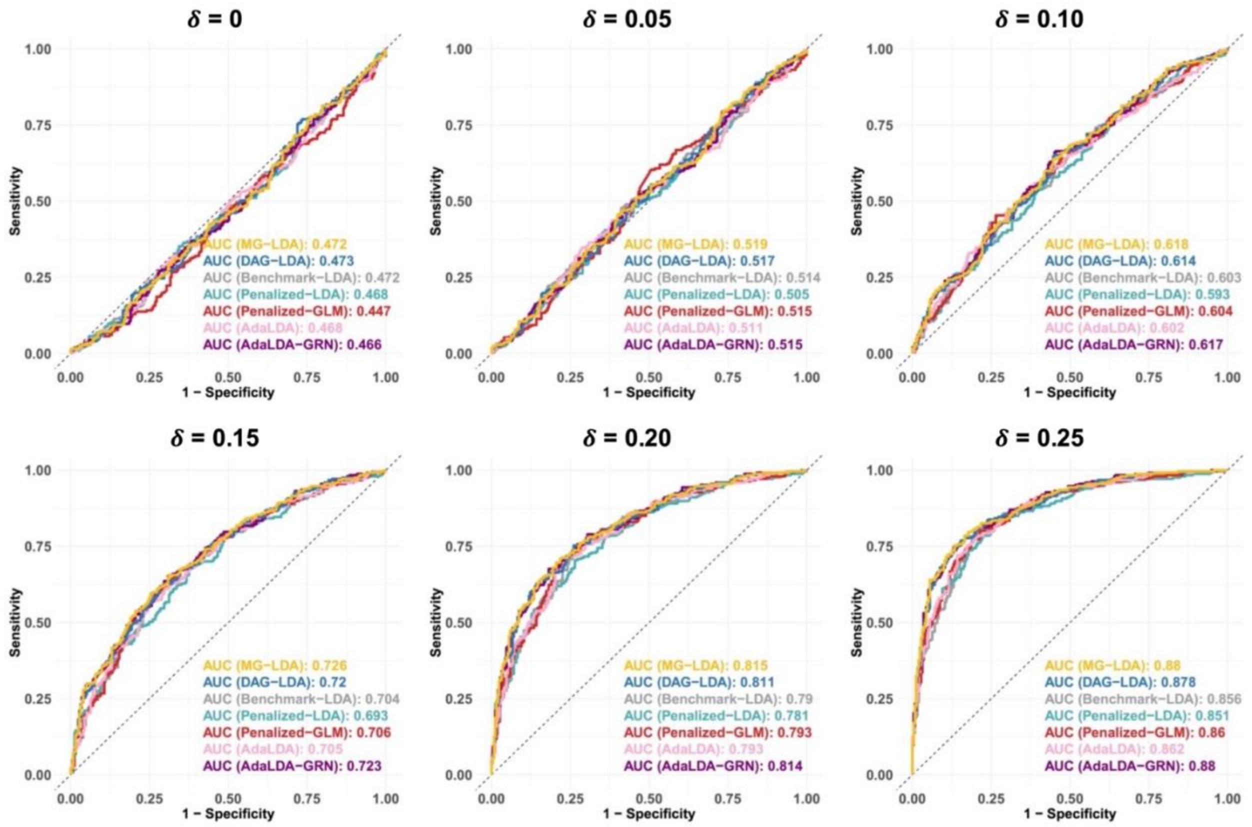


**Figure S4: Simulation results showing the classification performance of various LDA methodologies assuming $n_1 = n_2 = 50$, $p = 20$, and $p_0 = 0.4p$ under various settings of the mean difference between two groups $\delta$.** We consider MG-LDA and DAG-LDA: the proposed graph-informed LDA algorithm utilizing complete graph information (excluding self-loops) and the largest sub-DAG, respectively; Benchmark LDA: the standard Fisher's LDA; Penalized LDA, Penalized GLM, AdaLDA, and AdaLDA-GRN, which incorporates the GRN-informed covariance matrix generated by our GRN-informed method into the AdaLDA framework.

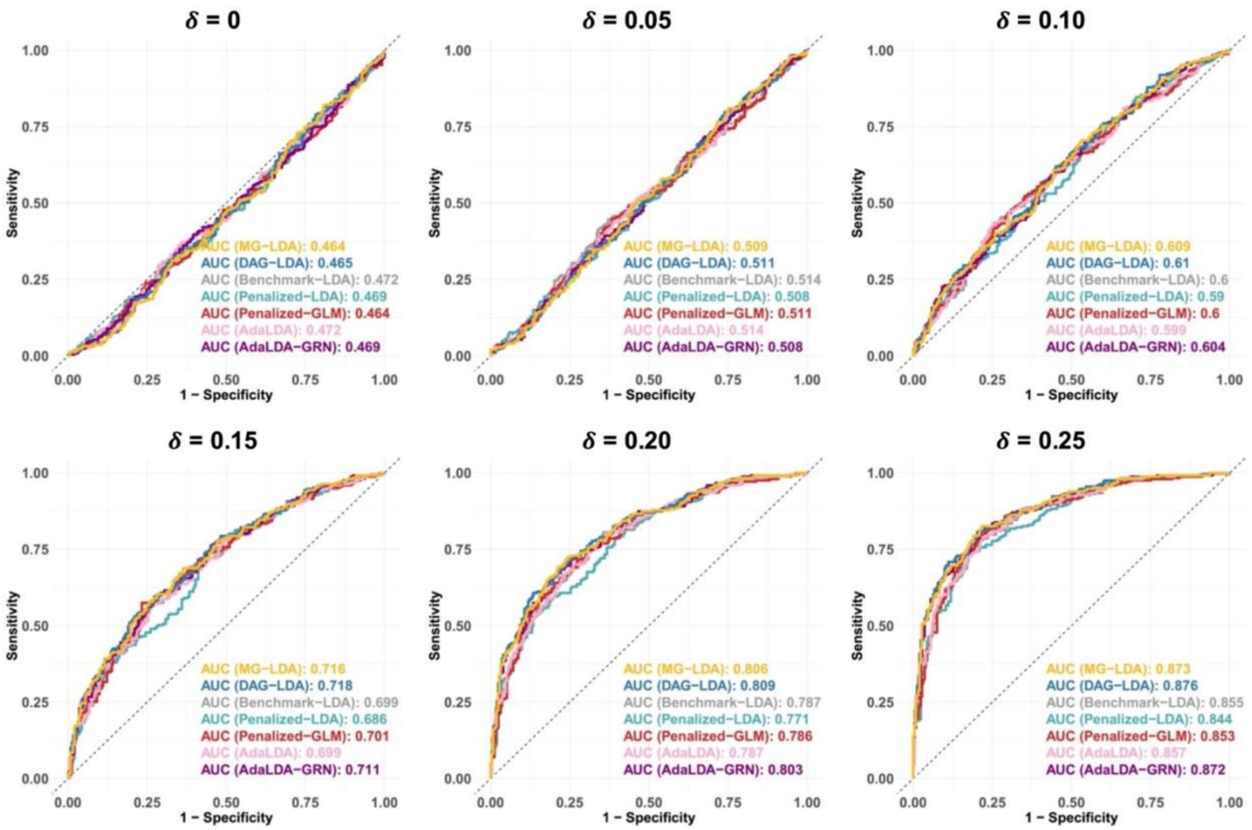


**Figure S5: Simulation results showing the classification performance of various LDA methodologies assuming $n_1 = n_2 = 50$, $p = 20$, and $p_0 = 0.8p$ under various settings of the mean difference between two groups $\delta$.** We consider MG-LDA and DAG-LDA: the proposed graph-informed LDA algorithm utilizing complete graph information (excluding self-loops) and the largest sub-DAG, respectively; Benchmark LDA: the standard Fisher's LDA; Penalized LDA, Penalized GLM, AdaLDA, and AdaLDA-GRN, which incorporates the GRN-informed covariance matrix generated by our GRN-informed method into the AdaLDA framework.

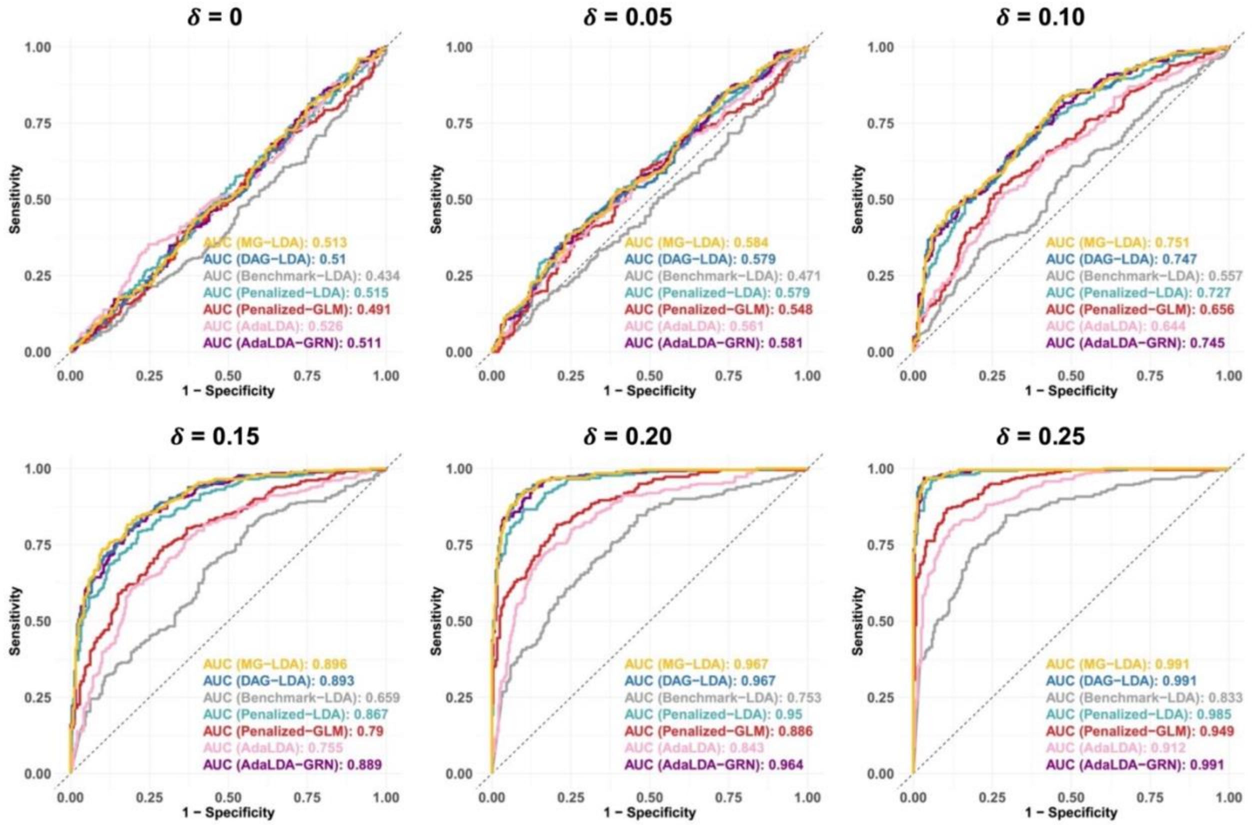


**Figure S6: Simulation results showing the classification performance of various LDA methodologies assuming $n_1 = n_2 = 50$, $p = 40$, and $p_0 = 0.8p$ under various settings of the mean difference between two groups $\delta$.** We consider MG-LDA and DAG-LDA: the proposed graph-informed LDA algorithm utilizing complete graph information (excluding self-loops) and the largest sub-DAG, respectively; Benchmark LDA: the standard Fisher's LDA; Penalized LDA, Penalized GLM, AdaLDA, and AdaLDA-GRN, which incorporates the GRN-informed covariance matrix generated by our GRN-informed method into the AdaLDA framework.

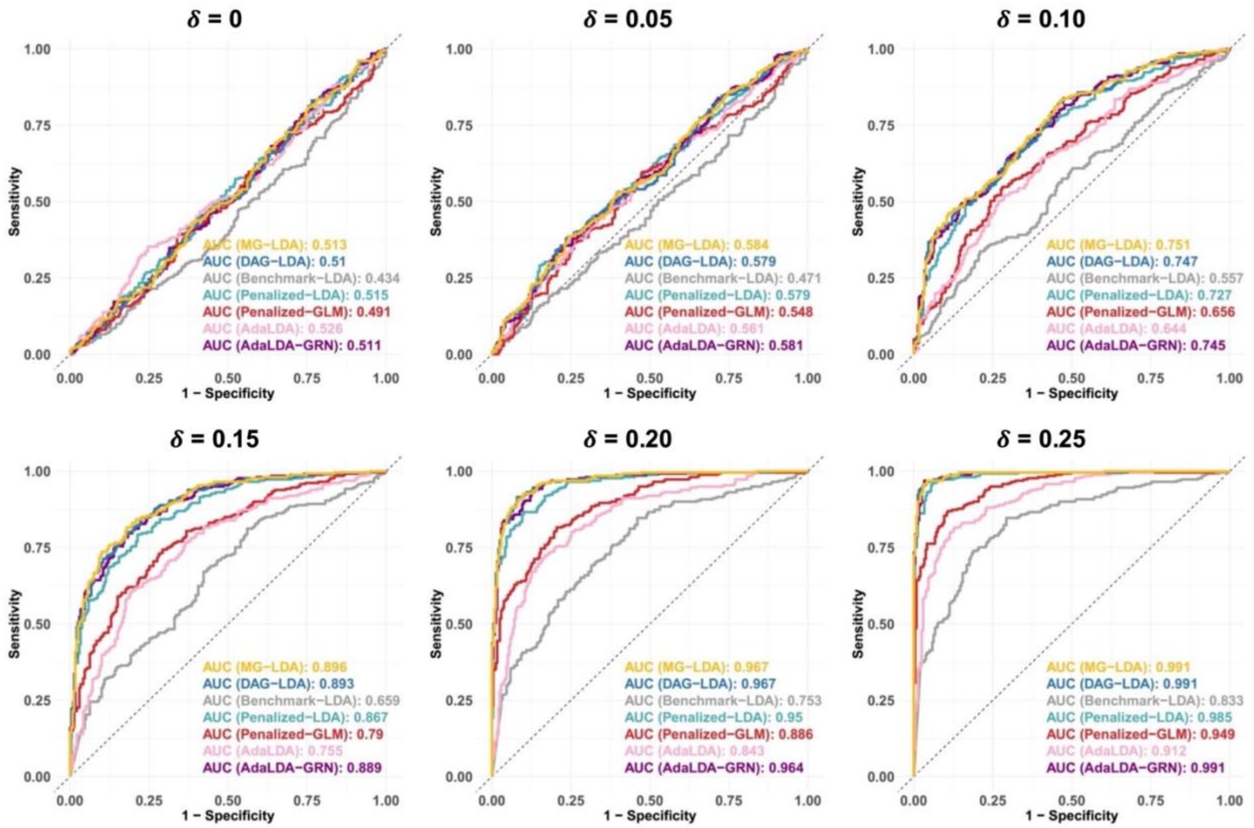


**Figure S7: Simulation results showing the classification performance of various LDA methodologies assuming $n_1 = n_2 = 100$, $p = 100$, and $p_0 = 0.4p$ under various settings of the mean difference between two groups $\delta$.** We consider MG-LDA and DAG-LDA: the proposed graph-informed LDA algorithm utilizing complete graph information (excluding self-loops) and the largest sub-DAG, respectively; Benchmark LDA: the standard Fisher's LDA; Penalized LDA, Penalized GLM, AdaLDA, and AdaLDA-GRN, which incorporates the GRN-informed covariance matrix generated by our GRN-informed method into the AdaLDA framework.

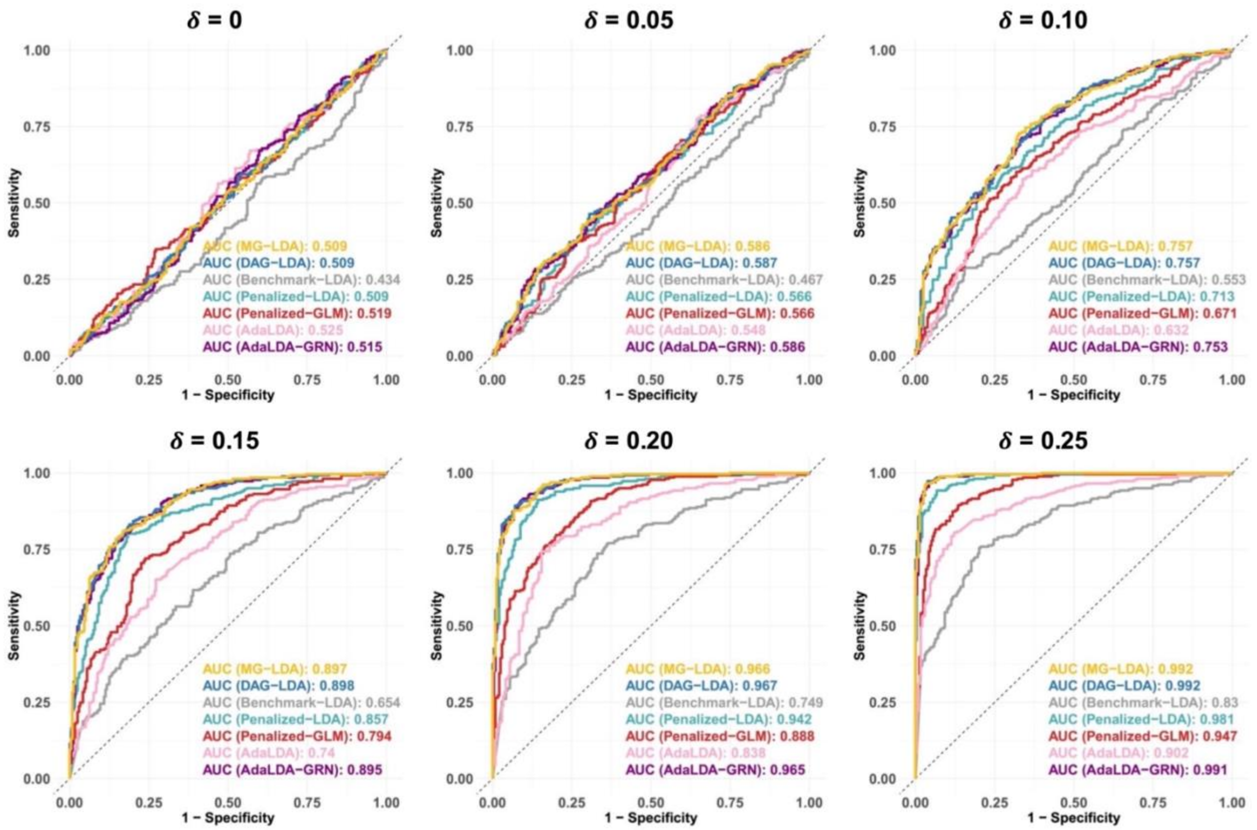


**Figure S8: Simulation results showing the classification performance of various LDA methodologies assuming $n_1 = n_2 = 100$, $p = 100$, and $p_0 = 0.8p$ under various settings of the mean difference between two groups $\delta$.** We consider MG-LDA and DAG-LDA: the proposed graph-informed LDA algorithm utilizing complete graph information (excluding self-loops) and the largest sub-DAG, respectively; Benchmark LDA: the standard Fisher's LDA; Penalized LDA, Penalized GLM, AdaLDA, and AdaLDA-GRN, which incorporates the GRN-informed covariance matrix generated by our GRN-informed method into the AdaLDA framework.

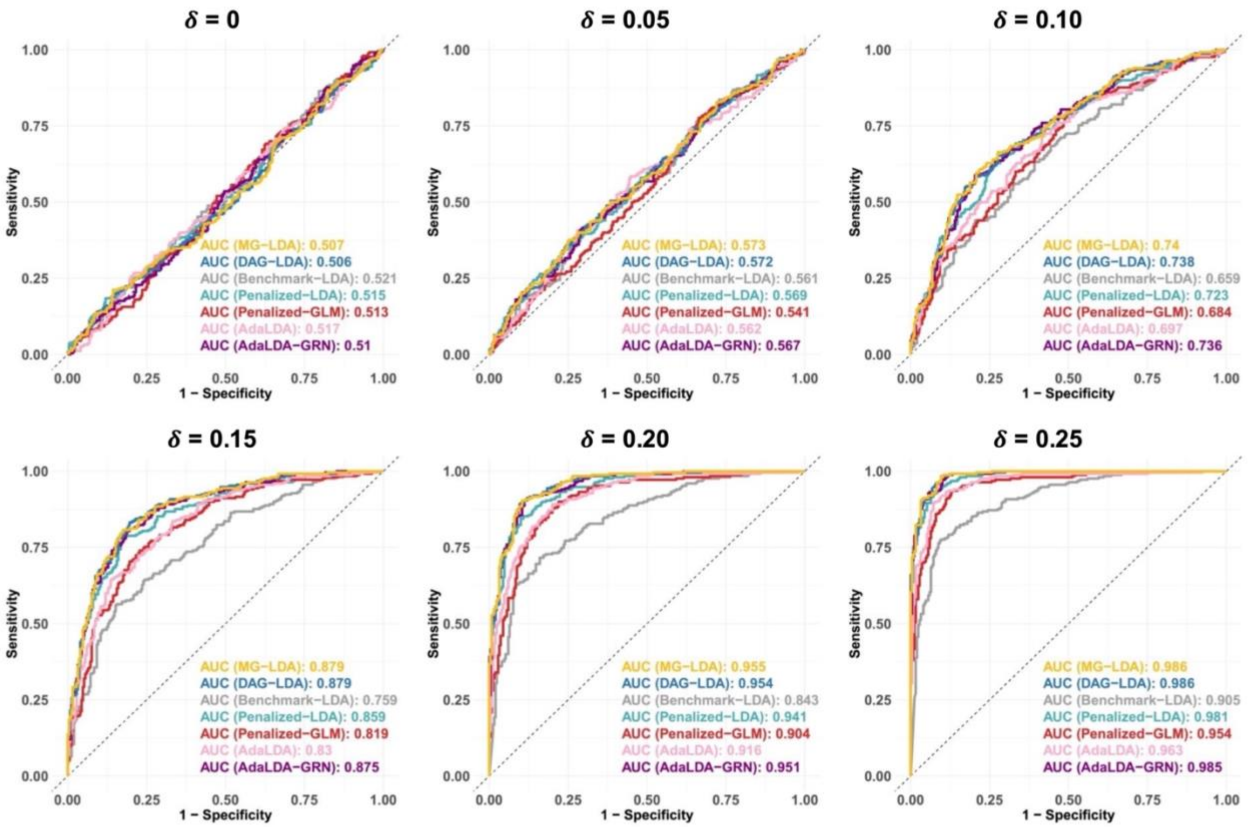


**Figure S9: Simulation results showing the classification performance of various LDA methodologies assuming $n_1 = n_2 = 100$, $p = 300$, and $p_0 = 0.4p$ under various settings of the mean difference between two groups $\delta$.** We consider MG-LDA and DAG-LDA: the proposed graph-informed LDA algorithm utilizing complete graph information (excluding self-loops) and the largest sub-DAG, respectively; Benchmark LDA: the standard Fisher's LDA; Penalized LDA, Penalized GLM, AdaLDA, and AdaLDA-GRN, which incorporates the GRN-informed covariance matrix generated by our GRN-informed method into the AdaLDA framework.

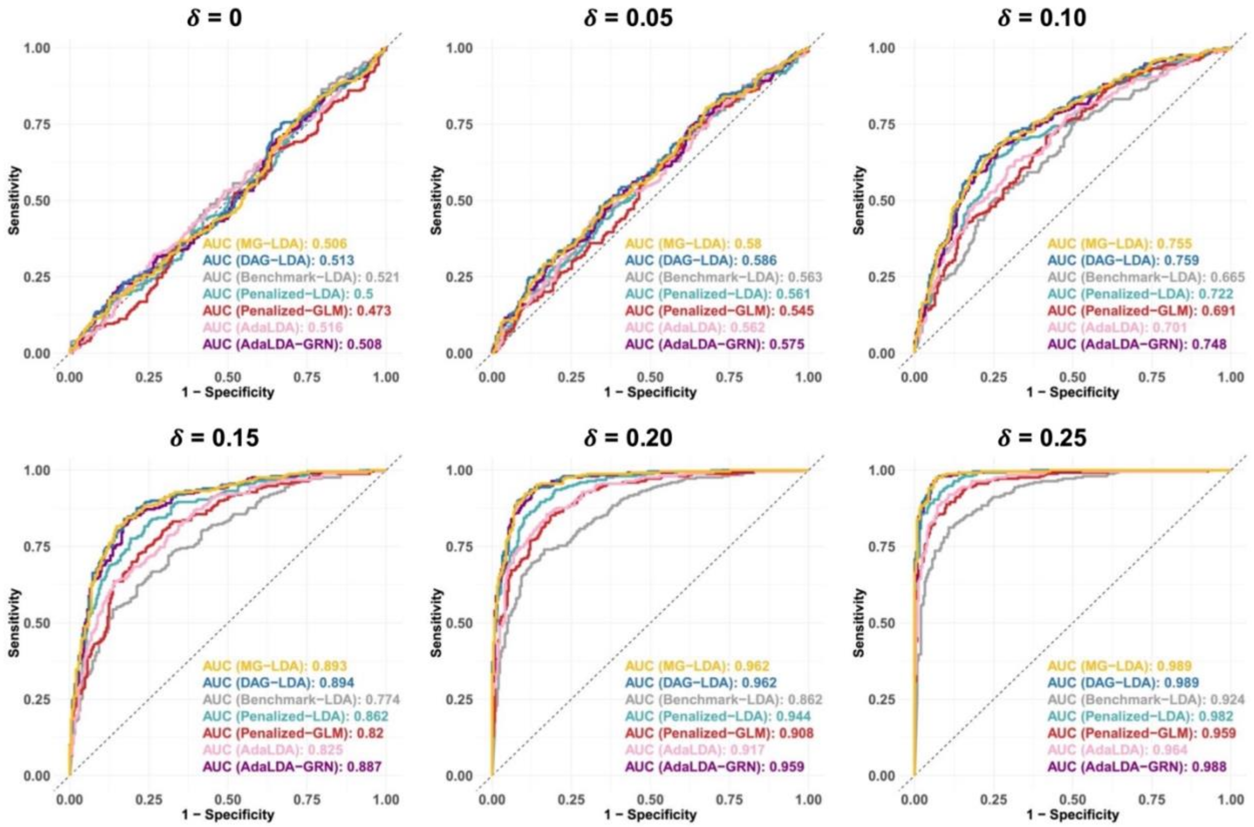


**Figure S10: Simulation results showing the classification performance of various LDA methodologies assuming $n_1 = n_2 = 100$, $p = 300$, and $p_0 = 0.8p$ under various settings of the mean difference between two groups $\delta$.** We consider MG-LDA and DAG-LDA: the proposed graph-informed LDA algorithm utilizing complete graph information (excluding self-loops) and the largest sub-DAG, respectively; Benchmark LDA: the standard Fisher's LDA; Penalized LDA, Penalized GLM, AdaLDA, and AdaLDA-GRN, which incorporates the GRN-informed covariance matrix generated by our GRN-informed method into the AdaLDA framework.
